# Predator type and relative risk affects the repeatability of nest defense in a songbird

**DOI:** 10.1101/2025.05.30.657090

**Authors:** Zoë M. Swanson, Chandler E. G. Carr, Josephina H. Fornara, Ross C. Eggleston, Dustin G. Reichard

## Abstract

Interindividual differences in nest defense towards a single predator type have been shown to be repeatable in multiple species, supporting the presence of personality. Here, we assessed if the nest defense of female northern house wrens (*Troglodytes aedon*) was repeatable across different types of predators, which is predicted by personality and largely untested. Over three years, we placed a decoy of a common nest predator, the eastern rat snake (*Pantherophis alleghaniensis*), on top of nest boxes and measured female behavior. Each season, we also presented a second stimulus of varying threat level, including an eastern chipmunk decoy (*Tamias striatus*), a taxidermied Cooper’s hawk (*Accipiter cooperii*), and a novel object of unknown threat. We measured repeatability between each pair of threats and compared the population-level response to the snake across years. Nest defense was significantly repeatable between the snake and chipmunk, which presented similar risks. It was also repeatable between the snake and novel object despite the population-level response to the object being significantly weaker. In contrast, nest defense was not repeatable between the snake and the hawk, which posed a significant threat to adult survival that may have disrupted the consistency of the female response. Finally, the average, population-level response to the snake did not detectably differ among years, indicating stability in this behavior despite high turnover in breeding adults. These results suggest the presence of personality in nest defense, but more research is needed to evaluate the effect of high-risk predators on the repeatability of this behavior.

**Significance Statement:** Differences among individuals in the strength of nest defense are often repeatable towards a specific predator, indicating the presence of personality. However, few studies have tested the repeatability of nest defense across different predators of varying threat levels. Personality predicts that weak/strong responders to one predator should also be weak/strong to another. We tested this prediction in female northern house wrens by placing decoys of different predators and a novel object on top of their nest boxes to measure nest defense. Nest defense was repeatable between two predators of similar risk and between a predator and a novel object of unknown risk. However, nest defense was not repeatable between a low and high-risk predator. The results suggest the presence of personality, but research is needed on the effects of high-risk predators that appear to disrupt behavioral consistency.

## Introduction

From circadian to circannual rhythms and affiliative to agonistic interactions, organisms repeatedly encounter the same or similar situations. Whether an individual responds to these repeated events with behavioral consistency or plasticity relates to their personality, defined here as consistent inter-individual differences (i.e., repeatability) in behavior across time and contexts (Réale et al. 2007; Roche et al. 2016). It is well established that animal personalities are common, and the field has pivoted away from documenting its occurrence toward investigating the ecological and evolutionary mechanisms that lead to its development and persistence (Laskowski et al. 2022). For example, the pace-of-life syndrome (POLS) hypothesis predicts that linkages between life history strategies, behavior, and physiological mechanisms could explain the persistence of inter-individual variation in behavior (Réale et al. 2010; Mathot and Frankenhuis 2018). Life history strategies fall upon a slow-fast continuum and fluctuations in selective forces through time and space might maintain variation rather than selecting for one optimal behavioral type (Dingemanse and Réale 2013; Mouchet et al. 2021).

Predation can have profound effects on the behavior and fitness of prey species leading to the development of proactive behaviors in anticipation of predators (e.g., changes in activity, reduced reproductive investment) and reactive behaviors when a predator is present (e.g., fight or flight; Caro 2005; Ghalambor et al. 2013; Gaynor et al. 2019). The appearance of a predator near eggs or offspring that cannot adequately defend themselves provides a context in which predation risk and personality can influence parental investment, a key component of life history theory (Trivers 1974; Clutton-Brock 1991; Ghalambor and Martin 2001). Parents face a tradeoff during nest defense between investing in their own survival (weaker defense) or their offspring’s survival (stronger defense). Individuals with a bolder, more aggressive personality that respond strongly to a low-risk predator may be more likely to protect their offspring at minimal cost to themselves (Montgomerie and Weatherhead 1988; Kleindorfer et al. 2003). However, those same individuals might also respond strongly to high-risk predators near their offspring and suffer an injury or death (but see Samelius and Alisauskas 2006). For instance, individual roach (*Rutilus rutilus*) with bolder personalities had a higher probability of being depredated than cautious individuals (Hulthén et al. 2017), indicating that predators can select against individuals that engage in riskier behaviors associated with a “fast” life history strategy (but see Smith and Blumstein 2008; Moiron et al. 2020). However, behavioral variation is usually maintained in free-living populations, which raises the question of why bolder personalities persist in the presence of predation?

One potential explanation is that bolder individuals can reduce their susceptibility to high-risk predators through threat assessment and behavioral plasticity. Rather than exhibiting the same response regardless of threat, successful individuals should exhibit plasticity in their nest defense based on the level of risk that each predator poses (Montgomerie and Weatherhead 1988; Clutton-Brock 1991; Kleindorfer et al 2005). Evidence from a variety of taxa indicates that antipredator behavior is plastic at the population level and varies with the level of risk, including during nest defense (Caro 2005; Lima 2009; Dassow et al. 2012). Studies of inter-individual variation in nest defense in birds have found that defense is often repeatable towards a single predator type (Kontiainen et al. 2009; Betini and Norris 2012; Burtka and Grindstaff 2013; Trnka et al. 2013; Clermont et al. 2019; Thys et al. 2019; Suckow et al. 2022; Suckow et al. 2022), but the repeatability of nest defense across different predator types, when the population-level response is expected to vary according to the perceived level of risk, remains largely untested (Salazar et al. 2021, 2023). It is important to note that the presence of plasticity does not indicate an absence of personality, a common misconception (Roche et al. 2016). Individuals can adjust their behavior across contexts while maintaining a similar response relative to others in their group. Repeatability estimates (*R*) are sensitive to group-level plasticity (Bell et al. 2009), and repeatability may be estimated as low and not significant when individuals react differently between sampling points, even if the inter-individual variation remains consistent. However, inclusion of a fixed effect, such as predator type, in the model allows for the calculation of an *adjusted* repeatability (*R_adj_*) estimate that controls for the variance caused by the fixed effect (Nakagawa and Schielzeth 2010). If the adjusted estimate is significant, then there is evidence that the behavior is repeatable despite group-level plasticity. In the case of nest defense, the population-level response is predicted to be weaker towards high-risk predators as all individuals reduce the strength of their response relative to a low-risk predator. Regardless of this population-level plasticity, individuals with bolder personalities should remain the strongest responders in the population across all predator types.

We investigated inter-individual variation in nest defense of female northern house wrens (*Troglodytes aedon*; hereafter “wrens”) in response to different threats at their nest box. In a previous study over two years (Eggleston & Fornara et al. 2025), we tested the nest defense of female wrens in response to a simulated eastern rat snake (*Pantherophis alleghaniensis*) and eastern chipmunk (*Tamias striatus*) in year one and the same rat snake decoy compared to a taxidermied Cooper’s hawk (*Accipiter cooperii*) in year two. We found that individual female wrens vary substantially in the strength of their nest defense but, at the population level, they respond similarly to nest predators that pose a low risk to adults (snake, chipmunk) and significantly less aggressively to a more dangerous predator (hawk; Eggleston & Fornara et al. 2025). Here, we report the results of a new analysis of repeatability in that existing two-year data set and an additional experiment in the same population to assess whether individuals exhibit personalities across varying threats. In the new experiment, which was conducted in the third consecutive breeding season, we compared female responses to the same snake decoy with the response to a novel object at the nest box. Unlike the threats posed by ecologically relevant predators, which the wrens can assess based on lived or evolutionary experience, the novel object’s threat level is unknown.

Our goals for this study were threefold. First, we compared female nest defense, at the population level, in response to the snake versus the novel object. Based on previous evidence for risk assessment in wrens and other species (Caro 2005; Lima 2009; Duré Ruiz et al. 2018; Eggleston & Fornara et al. 2025), we predicted that, on average, females would respond more strongly to the snake, which presented a high threat to offspring, but a low threat to adults, and weaker to the novel object, which presented an unknown threat. Second, we assessed the individual repeatability of female nest defense in three separate experiments that tested their response to simulated threats of varying risk. We predicted that nest defense would be significantly repeatable, which has been observed in other birds (see above), indicating that female wrens exhibit consistent, individual differences in their nest defense. We predicted that responses to threats of similar risk (snake v. chipmunk) would show limited plasticity while responses to threats of differing risk (snake v. hawk; snake v. novel object) would show high plasticity at the population level (Eggleston & Fornara et al. 2025). For individual differences to be repeatable when high plasticity is present, weak/strong responders to the snake should also be weak/strong responders to the hawk or dice, as evidenced by significant adjusted repeatability. Third, we analyzed whether the population-level response to the snake, which was included in each experiment, differed across three consecutive years. We predicted that the average, population-level response would be consistent among years, likely due to the continued maintenance of inter-individual behavioral variation in the population (Garamszegi et al. 2015; de Jong et al. 2021).

## Methods

### Study Sites and Species

Northern house wrens are cavity-nesting songbirds that defend their nests against a variety of predators, including woodpeckers, racoons, rodents, raptors, and snakes (Neill and Harper 1990; Dailey 2003; Millsap et al. 2013; Johnson 2020). We monitored 160 nest boxes for wren breeding activity between May 9 and August 25, 2022. The boxes were divided amongst six study sites in Delaware and Union counties in Ohio, USA (Eggleston & Fornara et al. 2025). Nest boxes were composed of wood and were mounted 1.75 meters high on a conduit pipe, which was coated with axle grease to limit predators. However, the boxes were located in edge habitat close enough to the surrounding vegetation that some ground-based predators could climb and access the boxes. The size of the entrance holes varied between sites due to the preferences of property owners, but all fell between 1.25 and 1.75 inches in diameter (Carr et al. 2024).

### Capture and Banding of Adult and Nestling House Wrens

We captured female wrens passively as they returned to their nest box in a 24 mm mist net. Each female was banded with a USFWS aluminum band and was assigned a unique color combination of three plastic bands for visual identification during behavioral trials. We took morphological measurements including the length of the wing, tail, tarsus, and bill, and mass. Feathers were measured with a ruler to the nearest mm, the tarsus and bill were measured with dial calipers to the nearest 0.1 mm, and mass was measured with a Pesola scale to the nearest 0.1 g. Any male wrens captured were also banded and measured for identification purposes, but most males were unbanded. We banded females at least 24 hours in advance of the first behavioral trial (X = 16.02 days, range = 1-60 days; 10 out of 43 females were banded within 7 days of their trial).

For each nest, we recorded the number of eggs laid, the number of offspring hatched, and the survival of those offspring to fledging. Due to logistical constraints, we were unable to check our boxes daily, and as a result, we were not always present to confirm every hatch day. However, wrens have a consistent incubation period and nestling growth rates that allow us to accurately estimate hatch dates and nestling age based on the onset of incubation, nestling mass, and feather development (Johnson 2020). After each behavioral trial, we counted the wren nestlings (X = 5.34, range = 4-8) and estimated their age. Once the nestlings reached eight days of age, they were banded with a USFWS aluminum band and morphological measurements were taken including tarsus and wing length and mass as described above. We combined these measurements with previous observations to confirm nestling ages (Brown et al. 2011).

### Experiment 1: Snake Decoy vs. Dice (Novel Object)

We followed a previously published protocol (Eggleston & Fornara et al. 2025) to conduct behavioral trials at 43 nest boxes containing nestlings from two to eight days old, between June 2 and July 29, 2022. A rubber model of a common wren predator, the eastern rat snake (*Pantherophis alleghaniensis*), and a pair of large white hanging car dice with black dots were used as stimuli. The car dice served as a novel object of unknown threat level. We chose the car dice because the dice were connected by a white string, which had a similar shape to the snake, but the dice themselves were unique with a furry texture. The white color of the dice should not have been interpreted as aposematic while standing out against the green nest boxes and foliage.

To initiate each trial, we waited for the female to return to the box before one observer flushed the female while placing the stimulus on the roof of the box. At the same time, a second observer placed a video camera two-to-three meters away from the nest box in an obscured location to record the trial. Once the stimulus was placed on the box, the observers retreated approximately 15 m from the nest box and monitored the female’s behavior for seven minutes. If the female did not return to the nest box within 3.5 minutes (half the trial), we reset the trial by removing the decoy only and waiting for the female to return to the nest box. Then, one observer flushed the female away and replaced the decoy to restart the seven-minute trial (23 of 86 trials; 26.7%).

During the trial, we recorded the number of flights over the stimulus (flyovers), number of direct contacts with the stimulus (hits), the closest approach to the stimulus, and time spent within five meters of the stimulus. The five-meter threshold was chosen because all boxes had perches within that distance despite heterogeneity among sites. We also noted whether females produced alarm calls during the trial (binary: yes or no). Treatment order was alternated from individual to individual to control for any effects of presentation order. Blind data collection was impossible because our study involved visual observations of free-living subjects.

After the trial concluded, one observer approached the nest box and removed the stimulus. The camera recorded for an additional 20 minutes to measure the amount of time it took the female to reenter the nest box. Six data points for return time (of 86; 7.0%) were lost due to camera malfunctions or video corruption. As a result, five females (of 43; 11.6%) that were missing one or both return time videos were excluded from the analysis for that behavior.

### Experiment 2 & 3: Snake Decoy v. Hawk or Chipmunk

Experiment 1 was replicated in the same wren population during the 2021 and 2020 breeding seasons with slightly different stimuli to compare female risk-taking behavior in the presence of different predators (Eggleston & Fornara et al. 2025). Here, we present a new analysis of these previously collected data to test for evidence of repeatability across antipredator contexts. In both experiments, we used the same eastern rat snake decoy but included a taxidermied juvenile Cooper’s hawk (*Accipiter cooperii*) decoy in Experiment 2 (2021) and a rubber eastern chipmunk (*Tamias striatus*) decoy in Experiment 3 (2020). The sampling methods and timing during the breeding season were the same as the Experiment 1 with a few key differences. A different observer recorded the female house wren’s behaviors in each experiment, but all were trained by the same person (DGR) to ensure consistent observations across years. In 2020, after the seven-minute trial concluded, the chipmunk model was reeled away from the nest box with a piece of fishing line to avoid the observer approaching the nest box. In 2021 and 2022, observers approached and removed the respective stimuli after the conclusion of the seven-minute trial. The females in the 2022 breeding season were captured in mist nets placed near their and banded in advance of the behavioral trials, while most females in 2020 and 2021 were unbanded, but males were captured and banded.

### Statistical Analyses

R version 4.4.2 (R Core Team 2024) was utilized for all the following analyses. For each experiment and the interannual comparison, we conducted a principal component analysis (PCA; packages: “stats,” “FactoMineR”, “factoextra”) to reduce the four major behaviors (flyovers, hits, time within 5 m, closest approach), which were correlated, into one composite response score. We used the “scale” function to standardize our variables before calculating the PCA. We retained only the principal components with eigenvalues greater than 1.0 and used the associated individual principal component scores (hereafter “PC scores”) for the subsequent analyses. Alarm calls were sampled as a binary response and analyzed separately. The time it took for the female to return to the nest box after the trial ended was also considered separately because it was sampled after the stimulus was removed.

For Experiment 1, we used separate linear mixed-effects models (LMMs, packages: “lme4,” “car”, and “lmerT-test”) that assumed a Gaussian distribution to test whether the PC scores and the time to return to the nest box differed between stimulus types (snake, dice). In each model, female behavior was the dependent variable and stimulus type, stimulus order, and their interaction were included as fixed effects. Day of the year was included as a covariate to evaluate seasonal changes in behavior, and female ID was included as a random effect to account for repeated sampling of the same individuals. To compare the production of alarm calls (recorded as presence/absence) between the stimulus types, we used the same model structure as above but shifted to a generalized linear mixed-effects model (GLMM) that assumed a binomial distribution. These analyses were not repeated for Experiments 2 and 3 because those results are reported elsewhere (see Eggleston & Fornara et al. 2025).

To assess the model assumptions, we tested model residuals for homoscedasticity with Levene’s tests (package: “car”) and normality using Shapiro-Wilk tests (package: “stats”), and we visually inspected plots of these and other assumptions using the “check_model” function of the “performance” package. The time to return to the nest box violated the assumptions of normality and homoscedasticity as the model residuals were right skewed. To resolve these issues, we square-root transformed the return time data and present those results here. The residuals for the PC score model met the assumption of normality, but failed Levene’s test despite appearing marginally homoscedastic when plotted. Linear mixed models are resilient to these violations, unless the sample size is prohibitively small or the distributions are bimodal (Schielzeth et al. 2020; Knief and Forstmeier 2021), neither of which occurs here.

For all three experiments, we assessed whether the PC scores, production of alarm calls, and time to return to the nest box were repeatable for individual females between stimulus types using the “rptR” package (version 0.9.22). The “rptGaussian” function was used for the PC scores and time to return, and “rptBinary” was used for the repeatability of alarm calls. For unadjusted repeatability (*R*) models, we included the behavioral response as the dependent variable, and the female ID as the grouping (i.e. random) factor. To account for the mean-level difference in response to decoys of different or unknown threat, we also modeled adjusted repeatability estimates (*R_adj_*) by including the type of stimulus (i.e., the identity of the decoy) as a fixed effect. All models were constructed with parametric bootstrapping (for rptGaussian, nboot = 10,000; for rptBinary, nboot = 1000) to produce 95% confidence intervals for the repeatability estimates to aid as an approximation of the robustness of the estimate. rptR uses Likelihood Ratio Tests to determine the significance of the repeatability estimate, with an alpha value of 0.05 being considered significant. Repeatability estimates greater than 0.4 were considered to be highly repeatable, while estimates between 0.20 and 0.40 were considered moderate, and estimates less than 0.20 were considered low (Carr et al. 2024). To estimate the proportion of variance in our adjusted models that were explained by the fixed effect (i.e., the variance removed), we processed the underlying lme4 model of each repeatability model through the “PartR2” package (version 0.9.2), at 1000 iterations of parametric bootstrapping to produce confidence intervals.

Because each female was sampled once with each stimulus type, we were unable to calculate any additional measurements of repeatability for the same decoy. Likewise, females were largely unbanded during the first two experiments, which made it impossible to test interannual repeatability. However, the replication of the snake stimulus across three experiments and years allowed us to test whether the population-level response varied through time. To test this possibility, we used generalized linear models (GLM; package: “stats”) assuming a Gaussian distribution with PC score or time to return to the nest box as the dependent variable and year as a fixed effect. Treatment order and day of the year were included as covariates to control for whether the snake was the first or second stimulus encountered and potential seasonal variation in behavior. The production of alarm calls was assessed using a GLM that assumed a binomial distribution. The PC score model met the assumptions of normality and homoscedasticity. The return time model met the assumption of homoscedasticity after a square-root transformation, but the residuals were not normal due to a persistent right skew. When a GLM yielded a significant fixed effect of year, pairwise comparisons were evaluated using the “emmeans” package.

## Results

### Principal Component Analysis (PCA)

For each PCA, all four behaviors loaded strongly onto the first component (PC1), which explained more than 50% of the total variation and was the only component with an eigenvalue above 1.0 (Table 1). In each relevant figure, the PC1 scores for Experiment 1 (Fig. 1A, 2A-B) are displayed as the opposite value for ease of interpretation. Thus, a higher PC1 score for each analysis is indicative of more hits, flyovers, and time spent within five meters of the stimulus, and a closer approach to the stimulus (i.e., more aggressive).

**Table 1.** PCA loading scores for each behavior measured.

| Principal Component<br>(% Variance<br>Explained) | Hits | Flyovers | Time Spent in<br>Five Meters | Closest Approach |
| --- | --- | --- | --- | --- |
| Exp. 1 PC1 (58.9%) | -0.437 | -0.527 | -0.550 | 0.478 |
| Exp. 2 PC1 (53.1%) | 0.441 | 0.481 | 0.489 | -0.579 |
| Exp. 3 PC1 (53.1%) | 0.383 | 0.456 | 0.592 | -0.543 |
| Interannual Snake PC1<br>(52.8%) | 0.402 | 0.477 | 0.584 | -0.520 |

**Figure 1.**
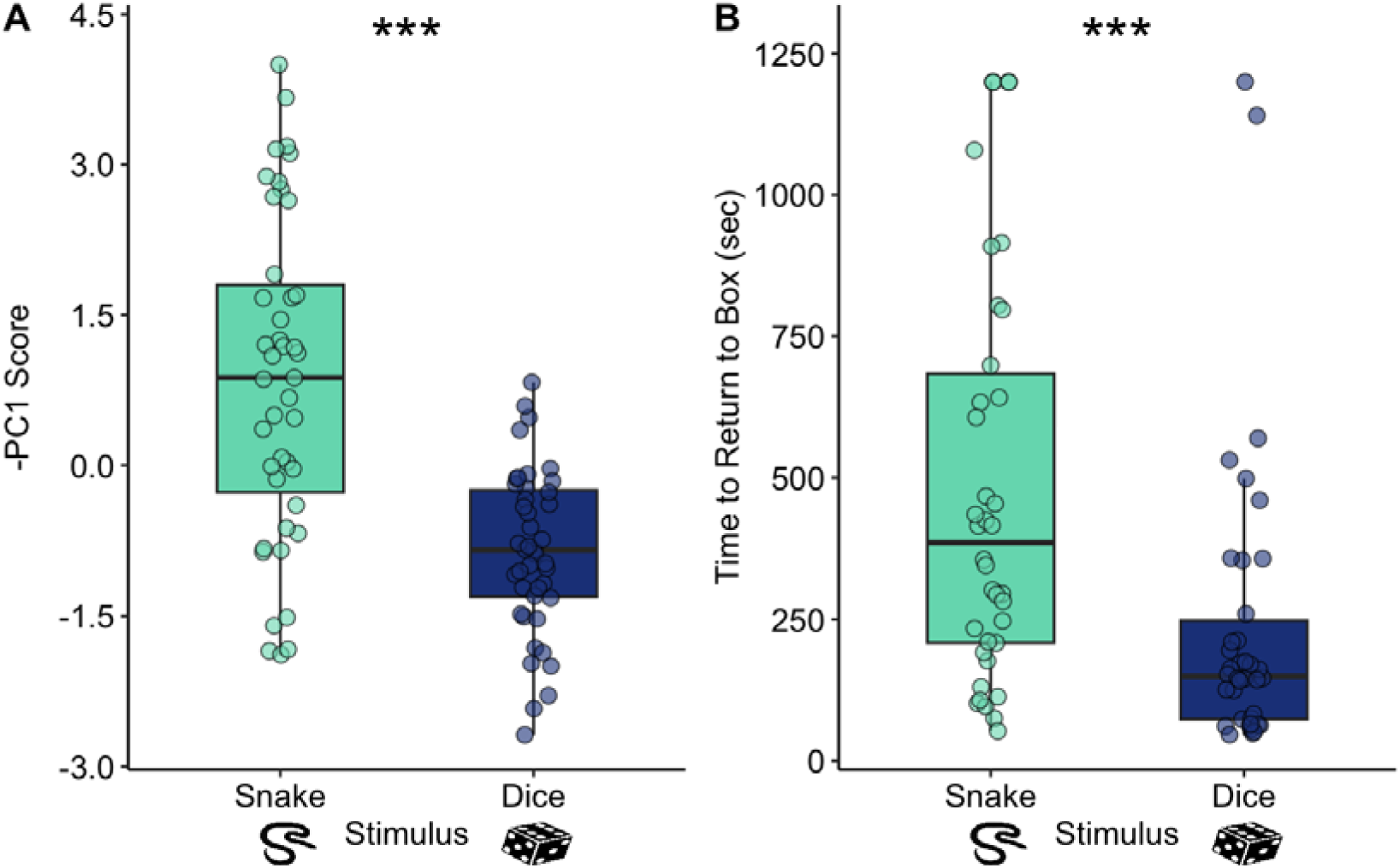
Female responses to the snake decoy and dice in Experiment 1. Higher PC1 scores in (A) represent more hits, flyovers, and time within 5 m of the stimulus, and a closer approach. Boxes show the interquartile range and median, and whiskers cover the range of data within 1.5 times the interquartile range. ***P<0.001.

**Figure 2.**
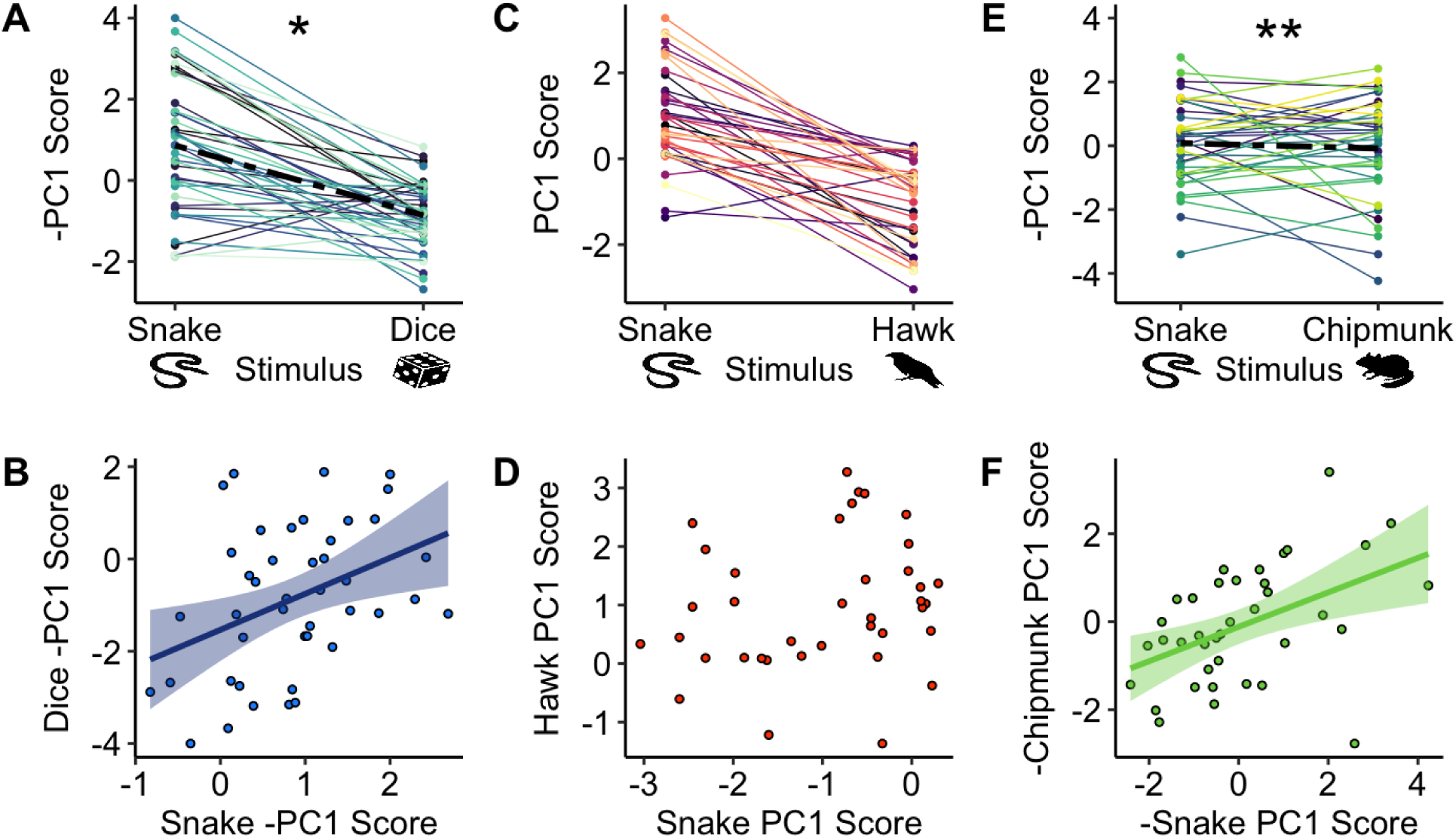
Reaction norms (A, C, E) and scatter plots (B, D, F) visualizing the repeatability of PC1 scores for Experiments 1 (A-B), 2 (C-D), and 3 (E-F). Higher PC1 scores represent more hits, flyovers, and time within 5 m of the stimulus, and a closer approach. PC scores were significantly repeatable in Experiment 1 (after adjusting for fixed effect; Radj=0.316, P=0.017) and Experiment 3 (unadjusted; R=0.458, P=0.002), but not in Experiment 2 (after adjusting for fixed effect; Radj=0.197, P=0.11). The thicker, broken line in (A) and (E) connects the mean responses for each stimulus. Line-of-best-fit in (B) and (F) is derived from a linear regression. *P<0.05. **P<0.01.

### Experiment 1: Snake Decoy vs. Dice (Novel Object)

PC1 scores were significantly different between treatments (*F_1,40.7_*=56.4, *P*<0.0001; Fig. 1A), where females responded more strongly to the snake than the novel object. There was a significant effect of date with weaker responses later in the breeding season (*F_1,40.0_*=6.90, *P*=0.01). The time to return to the nest box was also significantly different (*F_1,35.8_*=15.4, *P*<0.001; Fig. 1B), where females took more time to return in response to the snake than the novel object. Significantly more females produced alarm calls towards the snake decoy (36 of 42; 85.5%) than towards the novel object (25 of 42; 59.5%; *X^2^_1_*=5.28*, P*=0.02). We found no other significant effects of date, order, or the interaction of order and treatment on any of the behaviors (*P*>0.06 for all; Table S1).

### Individual Repeatability for all Experiments

PC1 scores were not significantly repeatable between the snake and the dice (*R*=0, SE=0.089 [95% CI=0-0.299], *P*>0.99), with the decoy explaining almost 32% of the behavioral response (*R²*=0.3165, 95% CI=0.1897-0.4479). After removing the variance associated with the decoy, the PC1 scores for Experiment 1 became moderately repeatable (*R_adj_*=0.32, SE=0.135 [95% CI=0.026-0.57], *P*=0.017, Figure 2). PC1 scores between the snake and the hawk decoys in Experiment 2 were not repeatable, even after accounting for the fixed effect (*R_adj_* =0.197, SE=0.139 [95% CI=0-0.482], *P*=0.11, Figure 2), which explained 47% (*R²*=0.4681, 95% CI=0.339-0.6052) of the variance. PC1 scores for Experiment 3 were highly repeatable between the snake and the chipmunk (*R*=0.463, SE=0.13 [95% CI=0.168-0.676], *P*=0.00158, Figure 2), with only 0.33% of the variance in response being explained by the decoy (*R²*= 0.0033, 95% CI=0-0.055).

Calls were not repeatable in Experiment 1 (*P*=0.482) but were highly repeatable in Experiment 2 (*R*=0.997, SE=0.017 [95% CI=0.946-0.999], *P*<0.001), with the decoy explaining none of the variance in the model (*R²*=0, 95% CI=0-0.0071). In Experiment 3, 89.5% (34/38) of individuals produced mobbing calls for both the chipmunk and the snake, indicating high consistency, but not repeatability, in this response. Time to return was not significant in any experiment, even after adjusting for the variance explained by the decoy (*P*>0.395 for all). See Table 2 for all unadjusted and adjusted individual repeatability estimates and *R^2^* estimates for decoy.

**Table 2.** Individual repeatability (R) and R² estimates for PC scores, Calls, and the Time to Return across years. Adjusted R values represent the proportion of variance explained by consistent individual differences after removing the variance associated with the Fixed effect. SE = Standard Error, 95% CI = 95% Confidence Interval around the estimate. Significant P-values are bolded.

| Experiment | Behavior | Unadjusted |  |  | Adjusted |  |  | Fixed |  |
| --- | --- | --- | --- | --- | --- | --- | --- | --- | --- |
| | | $R \pm SE$ | 95% CI | $P$-value | $R \pm SE$ | 95% CI | $P$-value | $R^2$ | 95% CI |
| 1<br>(Dice) | **PC1** | $0 \pm 0.089$ | 0-0.299 | >0.99 | **$0.316 \pm 0.135$** | **0.026-0.565** | **0.017** | 0.317 | 0.190-0.448 |
| | Calls | $0 \pm 0.086$ | 0-0.297 | >0.99 | $0.006 \pm 0.128$ | 0-0.365 | 0.48 | 0.083 | 0.006-0.248 |
| | Time to Return | $0 \pm 0.093$ | 0-0.31 | >0.99 | $0.123 \pm 0.128$ | 0-0.426 | 0.22 | 0.161 | 0.044-0.33 |
| 2<br>(Hawk) | PC1 | $0 \pm 0.094$ | 0-0.315 | >0.99 | $0.197 \pm 0.139$ | 0-0.482 | 0.11 | 0.468 | 0.339-0.605 |
|  | **Calls** | **$0.997 \pm 0.019$** | **0.933-0.998** | **<0.001** | **$0.997 \pm 0.017$** | **0.946-0.999** | **<0.001** | 0 | 0-0.007 |
| | Time to Return | $0 \pm 0.097$ | 0-0.319 | >0.99 | $0 \pm 0.10$ | 0-0.335 | 0.5 | 0.094 | 0.006-0.257 |
| 3<br>(Chipmunk) | **PC1** | **$0.463 \pm 0.13$** | **0.168-0.676** | **0.002** | **$0.458 \pm 0.131$** | **0.17-0.69** | **0.001** | 0.003 | 0-0.047 |
|  | Calls | - | - | - | - | - | - | - | - |
| | Time to Return | $0 \pm 0.103$ | 0-0.353 | >0.99 | $0 \pm 0.104$ | 0-0.344 | >0.99 | 0.005 | 0-0.09 |

### Interannual Comparison of Response to the Snake Decoy

PC1 scores did not differ detectably among the three years sampled (*X^2^* =3.99*, P*=0.14). There were also no significant differences in the number of females that produced alarm calls among years (*X^2^* =2.75*, P*=0.25). However, there was a significant difference in the time to return to the nest box (*X^2^* =14.9*, P*<0.001; Fig. 3B). Specifically, females took significantly longer to return to the nest in Experiment 1 (2022; *t*=-3.77, *P*<0.001) and in Experiment 2 (2021; *t*=-2.76, *P*=0.02) than in Experiment 3 (2020), but there was no significant difference between Experiments 1 and 2 (*t*=-0.99, *P*=0.59). We found no significant effects of treatment order or sampling date on any of these behaviors (*P*>0.07 in all cases, Table S2).

**Figure 3.**
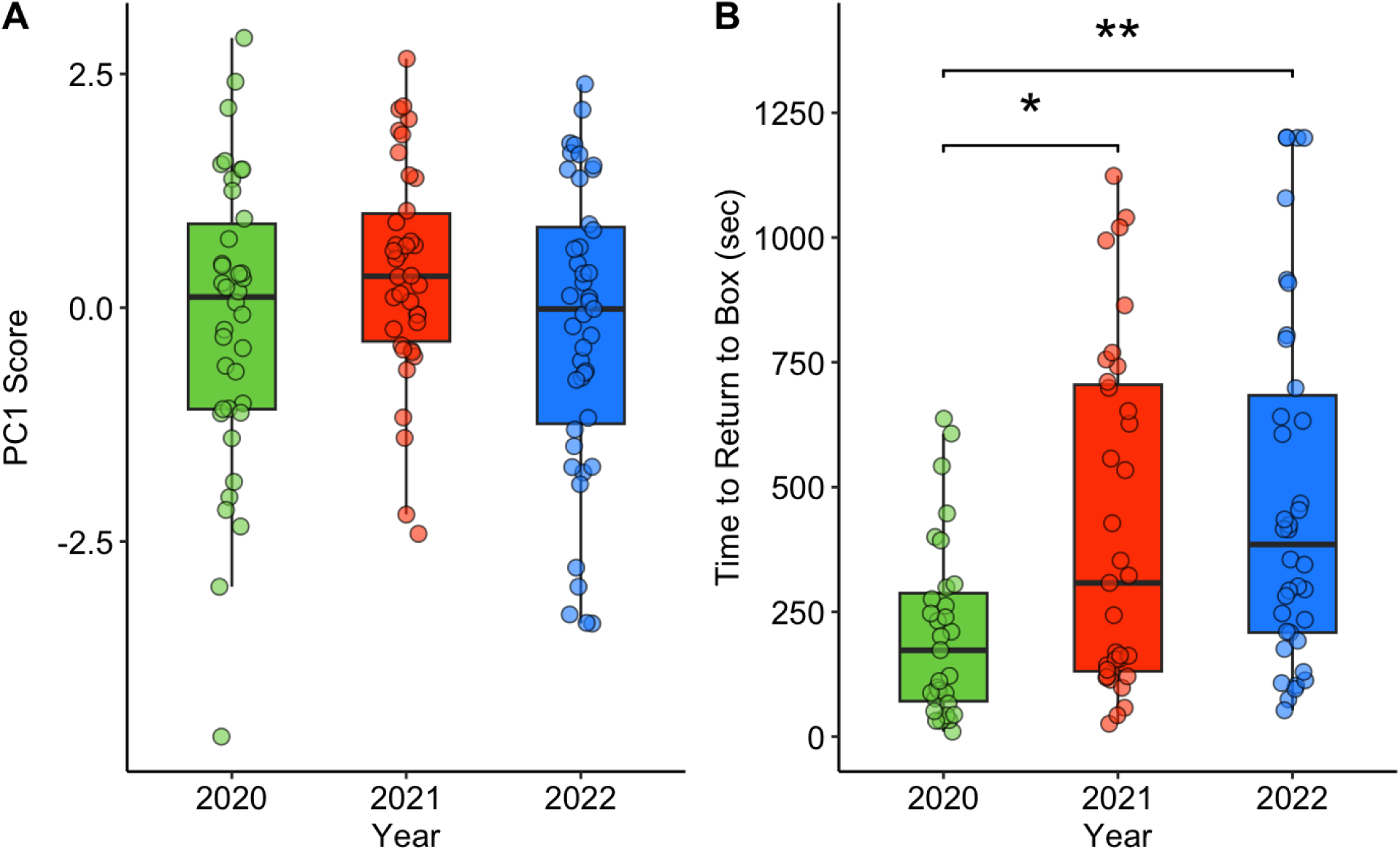
Interannual comparison of female responses to the snake decoy across three years. Higher PC1 scores in (A) represent more hits, flyovers, and time within 5 m of the stimulus, and a closer approach. 2020 = Experiment 3, 2021 = Experiment 2, 2022 = Experiment 1. Boxplots as in Figure 1. *P<0.05. **P<0.01.

## Discussion

In this study, we found that female northern house wrens responded significantly more aggressively to a simulated rat snake, which is a common nest predator, than to a novel object, which presented an unknown threat. Despite this population-level plasticity in nest defense, the female response was moderately repeatable between the snake and the novel object (*R_adj_* = 0.32), and highly repeatable between the snake and a simulated chipmunk (*R* = 0.46), another common nest predator. Both results support the presence of personality in nest defense. However, when comparing responses between the snake and a simulated Cooper’s hawk, a predator of both nests and adults (Millsap et al. 2013), we found no evidence of repeatability even when controlling for predator type. Collectively, these experiments indicate that females assess risk during nest defense and decrease their aggression when responding to threats of high (hawk; Eggleston & Fornara et al. 2025) or unknown (novel object) risk levels. The lack of repeatability in the female’s response to the snake and hawk did not match our predictions and suggests that the threat posed by the hawk disrupted the consistency of nest defense. Finally, we observed that the population’s average response to the snake did not differ detectably among three years, indicating that nest defense may remain stable through time despite high turnover among breeding adults and environmental heterogeneity between years.

### Risk Assessment

Prey species from across the taxonomic spectrum engage in risk assessment to modulate their antipredator behavior, often responding less aggressively to higher risk predators unless the benefits to their kin outweigh the risk to themselves (Montgomerie and Weatherhead 1988; Caro 2005; Lima 2009). Our results are consistent with previous studies of both northern and southern house wrens (*Troglodytes musculus*) that found evidence for risk assessment during nest defense (Duré Ruiz et al. 2018; Eggleston & Fornara et al. 2025), and we extended this analysis by comparing responses to a predator versus a novel object. In addition to responding more aggressively to the snake, significantly fewer females produced alarm calls in response to the novel object and females reentered the nest box significantly faster after responding to the novel object. A similar study in blue tits (*Cyanistes caeruleus*) also found that parents resumed provisioning faster after encountering a novel object near their nest, in this case a rubber ball, versus a simulated woodpecker, which frequently depredates nests (Mutzel et al. 2013).

The initial interaction with a novel object likely elicits caution from most individuals because the object’s risk to both themselves and their offspring is being assessed. Fewer individuals producing alarm calls may indicate that the novel object was not perceived as threatening to all female wrens, and a faster return to the nest box after the novel object was removed suggests that the object was considered a lower threat than the snake, which was an ecologically relevant nest predator (Mutzel et al. 2013; Johnson 2020). Parents that respond cautiously to novelty near their nests might avoid dangerous encounters, but more time spent away from the nest can be costly as offspring receive less food and heat (Dawson et al. 2005; Vrublevska et al. 2015; Dudeck et al. 2018; but see Mutzel et al. 2019). These relative costs will increase if the novel stimulus keeping parents away from their offspring carries no real threat, which removes any benefit of cautious behavior. Two studies in great tits (*Parus major*) supported these predictions as individuals that resumed provisioning young more quickly after the appearance of a novel object near their nest had higher nest success (Davidson et al. 2018) and female survival (Vrublevska et al. 2015).

### Repeatability of Nest Defense

The repeatability of nest defense towards a single predator has been established in multiple species (Kontiainen et al. 2009; Betini and Norris 2012; Burtka and Grindstaff 2013; Trnka et al. 2013; Clermont et al. 2019; Thys et al. 2019; Suckow et al. 2022; Suckow et al. 2022), but fewer studies have tested the repeatability of any behavior when responding to different types of predators at the nest. We found significant repeatability in nest defense (i.e., composite of hits, flyovers, approach measures) between predators of varying risk in two of three experiments. The snake and chipmunk were both predators of high risk to offspring and low risk to adults, making it unsurprising that individuals responded consistently to those predators. The novel object’s risk was unknown, and it elicited a weaker population-level response than the snake, but the individual responses were repeatable after controlling for predator type. Collectively, these results emphasize that although an individual’s behavior can be plastic across different contexts, the relative strength of the response among individuals can remain similar (i.e., high responders in context 1 are high responders in context 2). It is unclear what maintains persistent inter-individual differences in behavior, but multiple hypotheses are being investigated including genetic or epigenetic variation, state-dependent feedback mechanisms, and cues from the social environment (reviewed in Laskowski et al. 2022). Regardless, this study provides evidence that the expression of personalities is affected by predatory threat, but data from our remaining experiment suggest that nest defense is not repeatable across all threat types.

We did not detect any repeatability of nest defense when comparing the snake to the hawk, which is a predator of high risk to adults and low-to-moderate risk to nestlings (Millsap et al. 2013). Low values for repeatability (*R*) can either be caused by all individuals behaving the same (i.e., low inter-individual variation) or individuals lacking behavioral consistency between sampling events (i.e., high intra-individual variation; Roche et al. 2016). The reaction norms (Figure 2C) support the latter interpretation with individuals not responding consistently between the two predators. However, females were repeatable in whether they produced alarm calls to the snake and hawk, indicating that both predators were considered threatening. As an aerial predator, hawks are one of the largest threats to survival of adult songbirds, which likely explains the weaker population-level response to the hawk (Mahr et al. 2015; Eggleston & Fornara et al. 2025) and may have contributed to greater plasticity in nest defense among individuals. Whether intra-individual plasticity differed among individuals or across the three experiments is outside the scope of this study because females were only sampled once with each predator. Assessing individual variation in plasticity requires repeated sampling within and among different predator stimuli (Salazar et al. 2023), and this approach represents an important future direction for investigating how high-risk predators affect intra-individual variation in the plasticity of nest defense.

In a similar study that did not measure nest defense, Salazar and colleagues (2021) found that male blue tits were highly repeatable (*R*D=D0.51) in the latency to return to the nest after the departure of a predator when comparing responses to a human versus a taxidermic sparrowhawk (*Accipiter nisus*). In a subsequent study testing responses of blue tit pairs to a sparrowhawk, great spotted woodpecker (*Dendrocopos major*), and Eurasian blackbird (*Turdus merula*) as a non-predatory control, they again found evidence for repeatability across the different stimuli (*R*D=D0.18) despite population-level plasticity in the response to each threat (Salazar et al. 2023). In contrast, we found no evidence that the latency to return to the nest box was significantly repeatable in any of the three experiments included in this study, suggesting that this result may not be consistent across species. One important difference between these studies is that blue tits were sampled later in the nestling stage (days 10-12 post-hatch; Salazar et al. 2021, 2023) while wrens were sampled earlier (days 2-8 post-hatch) when the young are more dependent on female brooding for thermoregulation. The need to balance managing both the nest’s microclimate and nestling feeding might have disrupted the consistency of this behavior in females. Supporting this interpretation, female great tits that were incubating eggs, and thus not provisioning young, were moderately repeatable (*R*D=D0.25) in their latency to return to the nest when responding to a novel object versus predator eyes (Davidson et al. 2018). More sampling is needed across the nesting cycle in multiple species to fully disentangle the relationship between life history stage, predatory threats, and the inter-individual repeatability in the latency to return to the nest and other antipredator behavior.

### Interannual Variation in Nest Defense

The average, population-level nest defense and the number of females that produced alarm calls in response to the snake did not differ detectably among three years. This result is consistent with the frequent observation that behavioral variation is usually maintained over time as populations do not converge on one optimal behavioral type. Individual nest defense can be repeatable both within and among years (Burtka and Grindstaff 2013; de Jong et al. 2021), which could explain population-level consistency through time if many of the same individuals or their offspring are present. However, this explanation is unlikely to be supported in wrens, which are short-lived with a low return rate for adults (∼10-20%) and an even lower return rate for nestlings (<1%) in our population. These observations suggest that individuals from a variety of behavioral types are recruited into the population each year to maintain this population-level consistency. More generally, evidence from several long-term studies has indicated that selection on behavior fluctuates in both strength and direction across time and space, which provides one explanation for how behavioral variation and thus, population averages could be maintained (Dingemanse et al. 2004; Boon et al. 2007; Le Cœur et al. 2015; Mouchet et al. 2021).

In contrast to the nest defense results, we observed that the latency to return to the nest was significantly higher in the 2021 and 2022 breeding seasons, which can most likely be attributed to a change in our experimental protocol. In 2020, the snake decoy was pulled away from the nest box at the end of the trial with fishing line by humans sitting ∼15 m away, while in 2021 and 2022 a human walked to the nest box and quickly removed the stimulus. There was no detectable difference in latency to return between 2021 and 2022 when the removal by a human was replicated. These results are consistent with a similar study in great tits that found a longer latency to resume nestling feeding after a human disturbance at the nest box versus a human disturbance 20 m away (Mutzel et al. 2019). An individual’s willingness to approach a human threatening its kin or social group (Barnett et al. 2012; Vincze et al. 2019) and how long it takes to flee from an approaching human (i.e., flight initiation distance; Tätte et al. 2018; Grim et al. 2024) are common assays of boldness, which emphasizes the perceived threat elicited by humans. It is important for studies of antipredator behavior to consider how the presence of researchers can disrupt normal behavioral patterns and use techniques that minimize those effects.

## Acknowledgements

We thank Ohio Wesleyan University, the Big Walnut School System, and the Davis, Fink, and Koban families for access to facilities and field sites. We thank Ali Amer for field assistance and Lisa Tabak for help with equipment and other support. Animal images in Figures 1 2 were downloaded from PhyloPic.org and used in accordance with the <u>Creative Commons 3.0 License</u>. Credit to Chloé Schmidt for the chipmunk and to Ignazio Avella for the snake. Die image in Figures 1 and 2 was downloaded from commons.wikimedia.org and used in accordance with the <u>CC0 License</u>. Finally, thank you to the wrens that made this work possible.

## Data Availability Statement

The data and code to support these findings are freely available in the Mendeley Data Repository at DOI: 10.17632/4x44jyx4sg.1 and DOI: 10.17632/zjb5ynz5n4.4.

## Author Contributions

Conceptualization: ZMS, CEGC, JHF, RCE, DGR; Data Curation: ZMS, CEGC, JHF, RCE, DGR; Formal Analysis: ZMS, CEGC, DGR; Funding Acquisition: DGR; Investigation: ZMS, CEGC, JHF, RCE, DGR; Methodology: ZMS, CEGC, JHF, RCE, DGR; Project Administration: DGR; Resources: DGR; Supervision: DGR; Validation: CEGC, DGR; Visualization: ZMS, DGR; Writing - original draft: ZMS, CEGC, DGR; Writing - review and editing: ZMS, CEGC, JHF, RCE, DGR

