## Supplementary Materials for "Predator type and relative risk affects the repeatability of nest defense in a songbird"

*Table S1. Analysis of model fixed effects and covariates for Experiment 1. *P<0.05*

|  | Day of the Year | | Order | | Interaction with Order | |
| --- | --- | --- | --- | --- | --- | --- |
| Behavior | *F* or *Χ^2^* | *P* | *F* or *Χ^2^* | *P* | *F* or *Χ^2^* | *P* |
| PC1 Scores | 6.90 | 0.01* | 0.04 | 0.85 | 0.43 | 0.52 |
| Latency to Return | 0.02 | 0.88 | 0.18 | 0.67 | 3.86 | 0.06 |
| Alarm Calls | 0.05 | 0.82 | 0.22 | 0.64 | 0.67 | 0.41 |

*Table S2. Analysis of model fixed effects and covariates for interannual comparison of female response to snake decoy. *P<0.05*

|  | Day of the Year | | Order | |
| --- | --- | --- | --- | --- |
| Behavior | *Χ^2^* | *P* | *Χ^2^* | *P* |
| PC1 Scores | 0.95 | 0.33 | 0.07 | 0.78 |
| Latency to Return | 0.52 | 0.47 | 0.11 | 0.74 |
| Alarm Calls | 3.20 | 0.07 | 0.06 | 0.80 |
